# Sodium channel Nav1.2-L1342P variant displaying complex biophysical properties renders hyperexcitability of cortical neurons derived from human iPSCs

**DOI:** 10.1101/2021.01.18.427192

**Authors:** Zhefu Que, Maria I. Olivero-Acosta, Jingliang Zhang, Muriel Eaton, William C. Skarnes, Yang Yang

## Abstract

With the wide adoption of whole-exome sequencing in children having seizures, an increasing number of *SCN2A* variants has been revealed as possible genetic causes of epilepsy. Voltage-gated sodium channel Nav1.2, encoded by gene *SCN2A*, is strongly expressed in the pyramidal excitatory neurons and supports action potential firing. One recurrent *SCN2A* variant is L1342P, which was identified in multiple patients with early-onset encephalopathy and intractable seizures. Our biophysical analysis and computational modeling predicted gain-of-function features of this epilepsy-associated Nav1.2 variant. However, the mechanism underlying L1342P mediated seizures and the pharmacogenetics of this variant in human neurons remain unknown. To understand the core phenotypes of the L1342P variant in human neurons, we took advantage of a reference human induced pluripotent stem cell (hiPSC) line, in which L1342P was engineered by CRISPR/Cas9 mediated genome-editing. Using patch-clamping and micro-electrode array (MEA) recording, we found that the cortical neurons derived from hiPSCs carrying heterozygous L1342P variant presented significantly increased intrinsic excitability, higher sodium current density, and enhanced bursting and synchronous network firing, showing clear hyperexcitability phenotypes. Interestingly, the L1342P neuronal culture displayed a degree of resistance to the anti-seizure medication (phenytoin), which likely recapitulated aspects of clinical observation of patients carrying the L1342P variant. In contrast, phrixotoxin-3 (PTx3), a Nav1.2 isoform-specific blocker, was able to potently alleviate spontaneous and chemical-induced hyperexcitability of neurons carrying the L1342P variant. Our results reveal a possible pathogenic underpinning of Nav1.2-L1342P mediated epileptic seizures, and demonstrate the utility of genome-edited hiPSCs as an *in vitro* platform to advance personalized phenotyping and drug discovery.

## Introduction

Epilepsy is a devastating neurological disease characterized by recurrent episodes of seizures, resulting from the hypersynchronous firing of hyperexcitable neurons that go awry.^1^ Affecting about 50 million people worldwide,^2^ its etiology can vary and can be caused by genetic variants.^3^ With the broad adoption of whole-exome sequencing in children with seizures, genetic variants in *SCN2A* have been increasingly identified, emerging as one of the leading genetic causes of epilepsies.^4, 5^ *SCN2A*, encoding for the α-subunit of the voltage-gated sodium channel Nav1.2, is strongly expressed in principal neurons of the central nervous system, including excitatory glutamatergic neurons of the cortex.^6–9^ *SCN2A* is responsible for neuronal action potential (AP) initiation, propagation, and backpropagation during different developmental stages.^10, 11^ Studies have found “hot spots” in *SCN2A*, where variants occur frequently.^12^ In particular, a recurring heterozygous variant (Nav1.2-L1342P) has been identified in five patients, sharing similar and distinct features, including early-onset epileptic encephalopathy, transient choreoathetotic movements, and hypersomnia, among other symptoms.^5, 13–16^ Most of these patients have intractable seizures and are resistant to current medical treatments, which severely impairs their quality of life.

The channel properties of the L1342P variant have been previously studied in Human embryonic kidney (HEK293T) cells, revealing profound alterations in gating properties and a mixed gain-loss of function phenotype.^17^ However, how the L1342P variant affects the function of human neurons is unknown. Technological advances have made the generation of neurons from human-induced pluripotent stem cells (hiPSCs) possible, enabling the study of pathological features of human neurons carrying disease-associated variants *in vitro*. While patient-derived hiPSCs are often used for disease modeling, the genetic background of each patient may contribute to the phenotypes independent of the specific genetic mutation of interest. Additionally, when a variant affects more than one patient, it may result in different clinical manifestations. Thus, it would be important to assess the core phenotypes that can be attributed to that variant of interest, which will inform further pharmacogenetic discovery of such recurring variants. To this end, we utilized a clustered regularly interspaced short palindromic repeats (CRISPR)/Cas9 mediated genome-editing approach to generate the heterozygous L1342P variant in a well-characterized reference hiPSC (KOLF2) line.^18^ The KOLF2 line has a publicly available genome-sequencing information, is highly accessible for cross-lab comparison, and has a broad differentiation capacity into different cell types including neurons.^19–21^

By directly comparing neurons carrying the Nav1.2-L1342P variant with their isogenic WT controls, here we report that the reference hiPSC based *in vitro* model clearly shows that L1342P variant results in hyperexcitability phenotypes in neurons, including strong burst firing. Moreover, our *in vitro* platform is able to recapitulate aspects of pharmaco-responsiveness of patients and determine the efficacy of novel compounds. Our result is likely to inform the development of personalized therapy for epilepsy patients carrying specific genetic variants of *SCN2A*.

## Materials and methods

### Transfection of HEK293 cells with *SCN2A* plasmid

The human embryonal kidney cells (HEK293-tsA201 type) were cultured in Dulbecco's Modified Eagle Medium (DMEM) media with 10% fetal bovine serum (FBS) and 1% penicillin-streptomycin antibiotics. After reaching 80% confluency, cells were transiently transfected with Lipojet reagent (LipoJet™ In Vitro Transfection Kit, SignaGen Lab. Cat # SL100468) to introduce a full-length TTX-resistant human *SCN2A* WT or L1342P variant (gifts from Dr. Stephen Waxman, Yale University), in piggyBac vector with a GFP-2A linker, similar with the previously published design.^22^ After 24 hours, cells were dissociated into single cells using 0.25% trypsin (Corning, Cat # 25-053-CI) and plated on Poly-D-Lysine coated coverslips (Corning, Cat # 354086) for whole-cell patch-clamp recording.

### CRISPR-Cas9 editing of hiPSCs

CRISPR-Cas9 editing of KOLF2-C1 hiPSCs (p9) to generate Nav1.2-L1342P heterozygous single-cell derived clones was performed largely as described.^19^ Briefly, cells were grown in a 5% CO_2_, 37°C incubator in StemFlex media (ThermoFisher, Cat # A3349401) on Synthemax-treated wells (Synthemax II; Corning, Cat # 3535) and dissociated to single cells with Accutase (Stemcell Technologies, Cat # 07920). In a volume of 0.1 mL Primary Cell Buffer (Lonza), 1.6 x 10^6^ KOLF2-C1 cells were nucleofected (Lonza Amaxa 4D nucleofector; program CA137) with 20 mg Cas9 protein (HiFi; IDT), 16 mg single guide RNA (The first 20 bases: 5’-ATCTATCATGAATGTACTTC…-3’; chemically-modified; Synthego), and 200 pmol of a 100-mer oligonucleotide repair template (5’-GTTGTAAATGCTCTTTTAGGAGCCATTCCATCTATCATGAATGTACTTCCGGTTTGTCTGATCTTTTGGCTAATATTCAGTATCATGGGAGTGAATCTCT-3’; desalted Ultramer; IDT) containing the L1342P single nucleotide variant. Cas9 ribonucleoprotein complexes were assembled *in vitro* for 30 minutes prior to nucleofection. The cells were seeded onto one well of a Synthemax-treated 6-well plate and in StemFlex media containing 1X RevitaCell (ThermoFisher, Cat # A2644501), 30 mM HDR Enhancer (IDT), and cultured at 32°C (cold shock) for two days. After 24 hours, the media was changed to remove RevitaCell. After 48 hours, the media was changed to StemFlex without HDR Enhancer and cultured to confluency at 37°C. Following the cloning of single-cell derived colonies, genomic DNA was isolated. The target region containing the L1342P SNV was amplified by PCR (0.6 kb amplicon; forward primer, 5’-GGAATTTGATCCCCAAGTGGTCTCT-3’; reverse primer, 5’-AATGAGAGCTTTGCACTCACTGTAG-3’) and subjected to Sanger sequencing (sequencing primer, 5’-TTGGAGCTACCAGAGTCTAG-3’).

### Generation of hiPSC-derived neurons

Cortical neurons were generated with three to four individual differentiations using up to four different hiPSC lines (two for each genotype) to account for intra-cell line variability. We adopted a modified Dual-SMAD inhibition-based method, which simultaneously inhibits both Activin/Nodal and the Bone Morphogenetic Protein (BMP) signaling pathways with small molecules to drive cells into neuronal fate.^23–25^ In detail, hiPSCs were grown on a Matrigel substrate (Corning, Cat # 354230) in StemFlex Medium (ThermoFisher, Cat # A3349401) until 80-90% confluency in a 5% CO_2_, 37°C incubator. Colonies were passaged with an EDTA-based dissociation solution, Versene (Thermofisher, Cat # 15040066), and seeded with 10 mM rock inhibitor (RevitaCell Supplement, Gibco, Cat # A2644501) for the initial 24 hours on ultra-low attachment 96-well plates (Corning, Cat # CLS3474-24EA) with a cell density of 12,000 cells per well to obtain evenly shaped spherical embryoid bodies (EB). EBs were generated in an EB formation medium containing neural induction medium (Stemcell Technologies, Cat # 05835) supplemented with 100 nM LDN-193189 (Sigma, Cat # SML0559) and 10 μM SB431542 (Tocris, Cat # 1614). After seven days, EBs were harvested and replated into rosettes to generate neural progenitor cells (NPCs). A neural rosette selection reagent (Stemcell Technologies, Cat # 05832) was used to lift the rosette monolayer and allow expansion to obtain NPCs. Once neural progenitors were formed, they were plated on Poly-L-Ornithine (PLO)-laminin-coated coverslips and differentiated for the first seven days with a formula containing: neurobasal medium (Gibco, Cat # 21103049), 1X B27 plus supplement (Gibco, Cat # A3582801), 1X Non-Essential Amino Acids Solution (NEAA; Gibco, Cat # 11140050), GlutaMAX (Gibco, Cat # 3505006). Then the medium was switched into maturation medium, and cells were continued culturing for the next 38 days or longer containing: Brainphys (Stemcell Technologies, Cat # 05790), 1X B27 plus supplement (Gibco, Cat # A3582801), 1X MEM Non-Essential Amino Acids Solution (NEAA; Gibco, Cat # 11140050), GlutaMAX (Gibco, Cat # 3505006), 100 μM Cyclic adenosine monophosphate (cAMP) (Santa Cruz Biotechnology, Cat # sc-201567A), 200 μM ascorbic acid, 10 ng/mL brain-derived neurotrophic factor (BDNF) (ProspecBio, Cat # CYT-207) and 10 ng/mL glial cell-derived neurotrophic factor (GDNF) (ProspecBio, Cat # CYT-305). Cell media was replaced every 2-3 days.

### Immunocytochemistry

Immunocytochemistry was used to characterize the hiPSCs and determine the fate and maturity of differentiated neurons. Cells were maintained on glass coverslips (Neuvitro, Cat # GG-12-Pre) or 24-well glass-bottom plates with #1.5 cover glass (Celvis, Cat # P24-1.5H-N) previously coated with PLO-Laminin. On the day of the experiment, samples were washed briefly in phosphate-buffered saline (PBS 1X; Corning, Cat # 21-040-CMX12) and fixed in 4% paraformaldehyde in PBS at room temperature for 15 minutes. Samples were rinsed with PBS three times (5 minutes for each) and permeabilized for 20 minutes with 0.3% Triton X-100 (pH 7.4). Samples were blocked with 5% bovine serum albumin (BSA; Sigma Cat # 9048) for 1 hour at room temperature (RT), then left to incubate with diluted primary antibodies in 1% BSA in a humidified chamber at 4°C overnight. The next day, samples were rinsed three times with PBS and subjected to fluorescent-dye conjugated Alexa-Fluor-based secondary antibodies diluted in 1% BSA for two hours at RT in the dark. After incubation, the secondary antibody solution was removed, and coverslips were washed three times with PBS (5 min for each) in the dark. For DAPI counterstain, VECTASHIELD^®^ antifade mounting medium with DAPI (Vector Laboratories Cat # H-1200) or a PBS-DAPI solution (Thermofisher, Cat# 62238) (1:10000) was used.

Mouse-anti Paired box protein Pax-6 (PAX6) (Invitrogen, Cat #21103049) (1:300) and rabbit-anti Forkhead Box G1 (FOXG1) (Abcam, Cat # ab18259) (1:300) were used to stain neural progenitors. Rabbit-anti-beta Tubulin III (Abcam, Cat # ab18207) (1:1000), mouse-anti Microtubule-associated protein 2 (MAP2) (Invitrogen, Cat # 13-1500) (1:1000), guinea pig-anti Synapsin1/2 (Synaptic Systems, Cat # 106044) (1:1000), rabbit-anti VGLUT1 (Synaptic Systems, Cat # 135 302) (1:1000) and CTIP2 (Abcam, Cat # ab18465) (1:300) were used to stain hiPSC-derived neurons. Secondaries for both neural progenitors and hiPSC-derived neurons were anti-rabbit or mouse conjugated with Alexa-488 or Alexa-555, anti-guinea pig Alexa-488 and anti-rat Alexa-647 (Invitrogen) (1:1000). hiPSCs and neuronal cell imaging were acquired with a Widefield Nikon Eclipse Ti2 microscope. Images were processed using Nikon Imaging Software Elements. Representative Z-stack images for NPCs were acquired on a Nikon A1RMP inverted confocal microscope.

### Computational modeling

Computational modeling was used to predict the excitability determined by neuronal spiking changes. The numerical simulations were run under the NEURON environment (Version 7.72, http://neuron.yale.edu) with a previously established model.^26, 27^ The kinetic parameters of Nav1.2 were modified using a previously reported method according to activation and inactivation variables derived from our experimental data.^28^ Other parameters, including neuron physics and ion channel distribution, were in accordance with previous literature. To model the gating kinetics of the L1342P variant, the activation was set to vShift = −6 (vShifit = 10 in WT, and vShift change has an impact on both activation and inactivation). The number of action potentials was counted during the 20 ms progressively increasing current-injection period, starting from −75 mV membrane potential. The phase-plane plots were generated using the first triggered action potential waveform and were superimposed for comparison.

### Electrophysiology

Whole-cell patch-clamp recordings were performed with an EPC10 amplifier and Patchmaster v2X90.3 software (HEKA Elektronik, Germany) coupled to an inverted microscope (NikonTi-2 Eclipse). For voltage-clamp recording, experiments were conducted at room temperature with thick-wall borosilicate glass pipettes (BF150-86-10) with open-tip resistances of 2-4 MΩ. The external solution contained the following (in mM): 140 NaCl, 3 KCl, 1 CaCl_2_, 1 MgCl_2_, 10 HEPES, and 20 dextrose, titrated with NaOH to pH 7.3. The pipette solution contained (in mM): 140 CsF, 10 NaCl, 1.1 EGTA, 10 HEPES, and 20 dextrose, titrated with CsOH to pH 7.3. The osmolarity was adjusted with dextrose to 320 mOsm for the extracellular solution and 310 mOsm for the pipette solution. P/N procedure was used to subtract the leak currents. The measurement of the activation of voltage-gated sodium channel was achieved by 10 ms voltage steps from − 80 mV to + 50 mV with 5 mV increment, with a holding potential of −100 mV. The trace was first fitted with Boltzmann IV (Origin, version 9.7) to obtain the V_rev_ (reverse potential), then the sodium conductance was calculated as G_Na_ = I_Na_ / (V_m_ − V_rev_), where I_Na_ was the peak current amplitude (relative to the holding potential of −100 mV) under each corresponding voltage step V_m_. For steady-state fast inactivation analysis, the value was determined by using 500 ms pre-pulses to potentials from −140 to −5 mV, followed by a depolarizing test pulse to +5 mV. The window current was calculated as a percentage ratio between the overlapping area and the total area (over the range between −70 and 0 mV) under the activation and inactivation curves.

For current-clamp recording, thick-wall borosilicate glass pipettes (Sutter instrument, BF150-86-10) with open-tip resistances of 4-8 MΩ were pulled, and the same recording system was used. The external solution contained (in mM) 140 NaCl, 3 KCl, 2 CaCl_2_, 2 MgCl_2_, 10 HEPES, 15 dextrose, titrated with NaOH to pH 7.3; the internal solution contained (in mM) 140 KCl, 3 Mg-ATP, 0.5 EGTA, 5 HEPES, and 20 dextrose, titrated with KOH to pH 7.3. Dextrose was added to bring osmolarity to 320 mOsm and 310 mOsm for the extracellular and internal solution, respectively. After the cells were matured for 50 days, 2-3 differentiated neurons from each coverslip were recorded. The pyramidal-shaped neurons with multipolar dendrites were selected for electrophysiological recording to ensure consistent sampling. Action potential (AP) properties were analyzed from the first triggered AP in response to a series of depolarizing steps. Rheobase was determined by the minimum current stimulus that can trigger the first AP. The voltage threshold of the spiking waveform was determined at the voltage where a drastic upstroke is observed. The AP amplitude was measured from the peak to the fixed membrane potential of −75 mV. The spike width was determined by measuring the width of the AP when the potential reaches half of the maximum amplitude. The current densities were obtained by normalizing the peak current amplitude to the capacitance. For repetitive firing analysis, the number of action potential firings was recorded under each step of graded current injection during the range from 0 to 125 pA in 5 pA increment.

### Micro-electrode array (MEA) recordings on hiPSC-derived neurons

Neurons carrying both the Nav1.2-WT and heterozygous Nav1.2-L1342P variant were derived (two to three individual preparations from each genotype) from the same protocol (described in the generation of hiPSC-derived neurons) and studied with a procedure as identical as possible to minimize procedure-related variabilities. After 20 days of maturation, neuronal populations were dissociated into single cells using Accutase (Stemcell Technology, Cat# 07922), and 100,000 cells in 10 μL suspension were plated on a pre-coated (Poly-L-Ornithine) 48-well plate Cytoview plates (Axion, Cat# m768-tMEA-48W) compatible with a Maestro MEA system (Axion Biosystems), with each well containing 16 electrodes. We carefully controlled the seeding density to make sure each well had equal amount of cells between genotypes. Cells were maintained in Brainphys (Stemcell Technology, Cat# 05790) medium supplemented with 100 μM cAMP, 10 ng/mL BDNF, 10 ng/mL GDNF, 200 μM ascorbic acid, and 1X B27 incubated in 37°C for 14 days, with half-volume medium changes 2-3 times per week. On the day of recording, MEA plates with neurons were stabilized for 5 minutes in the system (Maestro, Axion Biosystems) in 5% CO_2_ at 37 °C. A total of 200 seconds of recording period was taken for analysis. The threshold for determining the voltage spikes was set to greater than six standard deviations from background noise. Active electrodes were defined as >5 spikes per minute of recording, and wells with less than 25% active electrodes were not included in data analysis. The spike waveforms were continuously monitored to ensure the electrodes were intact and functioning correctly. Synchrony index, bursting frequency, bursting duration, and bursting intensity (spiking frequency within each bursting event) analysis were obtained using the manufacturer software (Neural Metrics Tool, version 2.4. Axion Biosystems). The network synchrony between channels was measured based on the synchrony of spike firing between electrode pairs throughout the whole well.^29^ A burst was defined as a collection of a minimum of five spikes, each separated by an inter-spike interval (ISI) of less than 100 ms.

The compound-testing was performed using the same 48-well plate during the culture time between 45-55 days. The plate was placed on the MEA system for 5 minutes, followed by a 100-sec recording to obtain the baseline measurement. Then, compounds were added to the MEA in a sterile condition, which was followed by another 5-minute equilibrium. Afterwards, a 100-sec recording was made to study the drug effect. Fold changes were calculated by comparing the compound treatment to baseline (for phenytoin and PTx3 baseline inhibition) or to kainic acid (KA) (for PTx3 inhibition on KA-induced firing). Drugs used were: kainic acid monohydrate (Sigma-Aldrich, Cat # K0250. 5 μM); phrixotoxin-3 (PTx3) (R&D Systems, Cat # 4914. 15 μM); phenytoin (Sigma-Aldrich, Cat # PHR1139. 15, 30, 40, 50 μM). Stock solutions of KA and PTx3 were prepared in autoclaved MilliQ water. Stock solutions of phenytoin were made in DMSO (final concentration of DMSO < 0.2%).

### Statistical analysis and criteria for neuron selection

Normality test was performed for all data. Data were analyzed using a parametric t-test, a nonparametric Mann-Whitney *U* test or Wilcoxon test (If data is not normally distributed), or a repeated-measures one-way or two-way ANOVA on Origin (OriginLab) and Prism (GraphPad Software). Data are reported as mean ± standard error of the mean (SEM). The significance threshold was set with a p-value less than 0.05, with p < 0.05 (*), p < 0.01 (**), and p < 0.001 (***).

### Data availability

The data that support the findings of this study are available from the corresponding author, upon reasonable request.

## Results

### L1342P variant in Nav1.2 displayed a larger window current indictive of a gain-of-function feature

The α-subunit of Nav1.2 comprises four homologous domains (I to IV), with each domain containing six transmembrane spanning regions (S1-S6). S1-S4 forms a voltage sensor domain (VSD), and the S5-S6 helix with an ion selectivity filter forms the pore domain to permit sodium influx (**Fig. 1A**). To visualize the location of the L1342P variant, we utilized a 3D human Nav1.2 structural model (PDB ID: 6J8E) (**Fig. 1B**).^30^ L1342P is positioned at the fifth segment (S5) on domain III, which is a highly conserved region.^17^ The fifth segment and neighboring locations are the hotspots for many seizure-related variants.^12^ L1342P is predicted to disturb interactions with the adjacent S4-S5 linker.^17^ The structural significance of the S5 region implies that the biophysical properties of Nav1.2 are likely to be profoundly disrupted by the L1342P variant.

**Figure 1.**
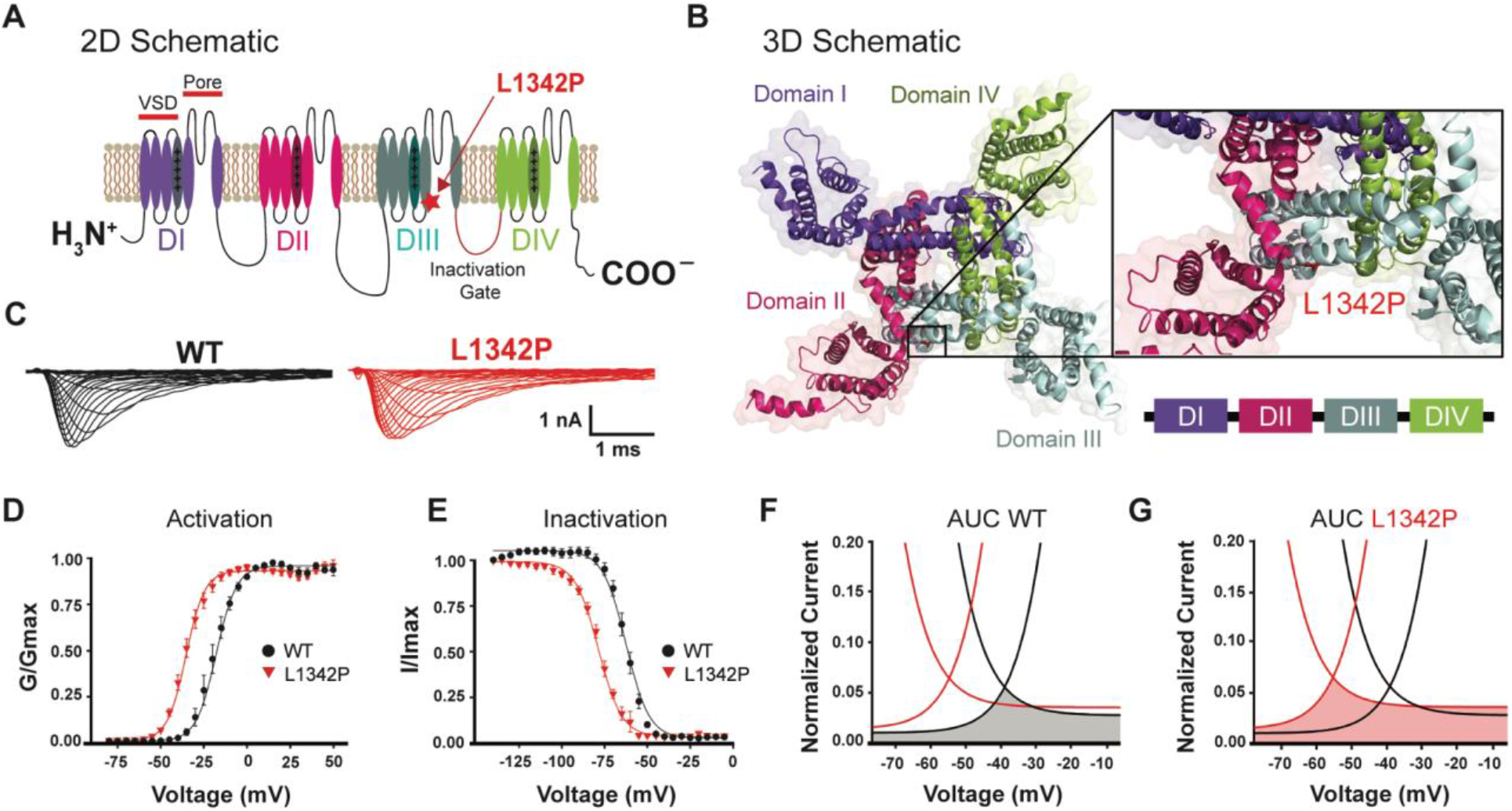
Structural modeling and biophysical properties of L1342P variant. (**A**) Schematic transmembrane topology of Nav1.2 sodium channel. Mutated residue L1342P is located at domain III, fifth segment, and is denoted as a red star. (**B**) Three-dimensional view of the structural model of Nav1.2 channel transmembrane domains. The location of the L1342P variant is at the S5 helix of the pore-forming segment. A zoomed-in view was shown on the right. (**C**) Representative traces for whole-cell sodium current from HEK cells expressing wild-type Nav1.2 (left) or Nav1.2-L1342P channel (right). (**D**) Voltage-dependent activation. The normalized conductance was plotted against the voltages in the test pulses, ranging from –75mV to 50 mV, then fitted with the Boltzmann equation. (**E**) Steady-state fast inactivation of wild-type and L1342P mutant channels. The relationship between the normalized current peak amplitude and prepulse potential was plotted and fitted with the Boltzmann function. (**F-G**) Illustrative presentation of window current analysis for WT (F) and L1342P (G). The shaded area depicts window current, which is the overlapping region under activation and inactivation curves.

To examine the effect of the L1342P variant on the biophysical properties of ion channels, we performed whole-cell voltage-clamp recordings on human embryonal kidney (HEK) cells. Two representative whole-cell current traces from cells expressing either WT or Nav1.2-L1342P mutant channels are shown, indicating that functional sodium currents can be recorded from both WT and Nav1.2-L1342P channels (**Fig. 1C)**. The L1342P exhibited a large, hyperpolarized shift of V_half_ in voltage-dependent activation compared to WT, suggesting a gain-of-function phenotype (**Fig. 1D**) (V_half_ value: WT: –19.5 ± 0.5 mV, n = 9; L1342P: –35.4 ± 0.5 mV, n = 11; p < 0.001, *student’s t-test*). Steady-state fast-inactivation was measured in response to a 500 ms depolarizing potential. L1342P markedly shifted channel fast-inactivation V_half_ by around 16 mV in a depolarizing direction (**Fig. 1E**) (V_half_ value: WT: –61.6 ± 0.1 mV, n = 6; L1342P: –77.8 ± 0.3 mV, n = 11; p < 0.001, *student’s t-test*), suggesting a loss-of-function attribute. Taking channel activation and fast-inactivation together, the L1342P variant resulted in a complex biophysical change at the channel level, showing both gain and loss-of-function phenotypes for different parameters. To further understand the overall effect of these biophysical changes, we calculated the window current. Window current is defined as the inward current that arises from partial activation and incomplete inactivation of the sodium channel, and the magnitude of which can be used to assess the net effect of gain- versus loss-of-function of a particular variant.^31, 32^ Our calculation revealed that the magnitude of the window current from L1342P is greater than that from WT (WT: INa/total = 5.8%; L1342P: INa/total = 7.0%) (**Fig. 1 F-G**), likely contributing to an overall gain-of-function phenotype of the L1342P variant.

### Single-compartmental simulation predicts a gain-of-function phenotype in a pyramidal neuron model with the Nav1.2-L1342P variant

To further study how the L1342P variant of Nav1.2 alters neuronal excitability, we used the NEURON simulation to model the firing properties of pyramidal neurons, in which Nav1.2 is strongly expressed.^6–9^ Modeled with the biophysical properties measured in our HEK293 cell study, we found that the neuron with Nav1.2-L1342P is notably more excitable. We define the minimal current inject to elicit the AP firing of neurons modeled with L1342P as 1X. This same 1X current injection, however, cannot trigger action potential firing in neurons modeled with WT Nav1.2 channel (**Fig. 2A)**. Number of APs from neurons modeled with Nav1.2-L1342P increases in response to elevated current injection. At each current injection level, neurons modeled with Nav1.2-L1342P had more action potential firing than neurons modeled with WT Nav1.2 channel over a broad range of current injections (**Fig. 2B**). To evaluate changes in single action potential waveforms, phase plane plots were constructed from the first derivative of the membrane potential (dV/dt) versus the membrane potential. We found a slightly lower action potential threshold in the L1342P neurons, with no detectable difference between WT and L1342P neurons in peak amplitude or maximum value of dV/dt (**Fig. 2C**). Collectively, our results from the computational modeling suggest an overall hyperexcitability phenotype in neurons carrying the Nav1.2-L1342P variant, which prompts us to test this prediction in human neuron-based models.

**Figure 2.**
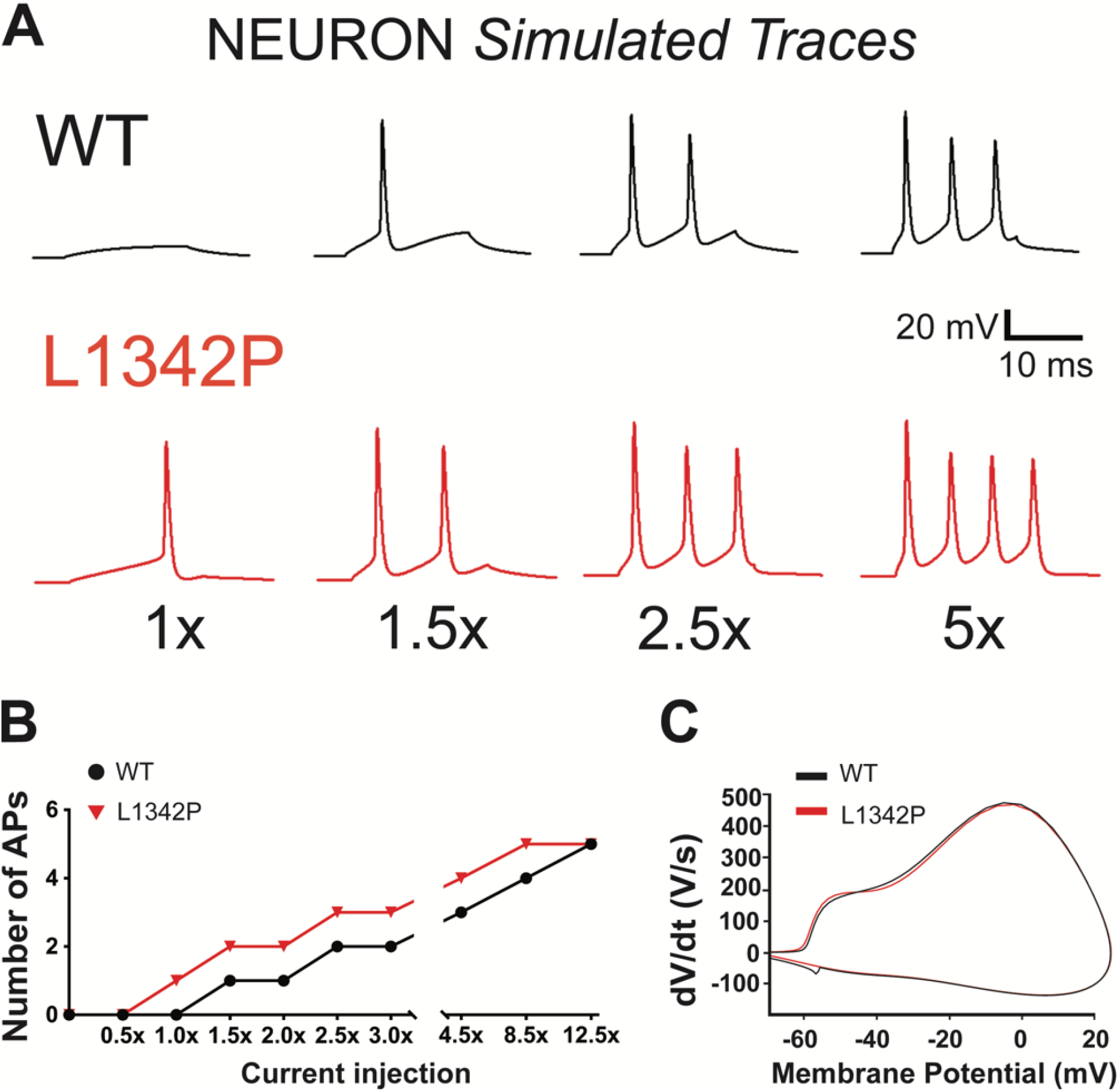
Single-compartmental simulation in a virtual neuron with Nav1.2-L1342P variant predicts gain-of-function. (**A**) Computational simulation predicated a higher firing frequency in the virtual neuron model with the L1342P variant. (**B**) The input-output relationship showed a significantly increased number of spikes in the range of current injections in L1342P variant. (**C**) Phase-plane plot comparison between virtual WT and L1342P neuron. The virtual L1342P neuron had a slightly hyperpolarized voltage threshold, while no significant differences were found in other parameters.

### hiPSCs carrying the L1342P variant can be differentiated into a cortical glutamatergic neuronal fate

Excitatory glutamatergic neuron in the cortex is a key cell type involved in seizures,^4, 28, 33^ and the methodology to reliably generate cortical glutamatergic neurons from hiPSC is well established.^34–36^ To understand how the Nav1.2-L1342P variant affects the function of human cortical neurons, we used CRISPR/Cas9 to recapitulate the sequence change of heterozygous Nav1.2-L1342P in a reference hiPSC (KOLF2) line (referred to L1342P thereafter).^19^ The Nav1.2-L1342P variant has been identified from multiple patients with both shared and distinct phenotypes.^5, 13–16^ Thus, a reference hiPSC line has the advantage of reducing the influence of the genetic background across individual patients and allowed us to assess the core impact of the Nav1.2-L1342P variant on neurons compared to isogenic controls.

After CRISPR/Cas9 mediated editing process, we successfully obtained both isogenic WT and heterozygous L1342P hiPSCs validated by sequencing (**Fig. 3A**), which can be differentiated into glutamatergic cortical neurons using an established protocol (**Fig. 3B**). Undifferentiated hiPSC colonies displayed normal and homogenous morphology with defined edges and low levels of spontaneous differentiation. They consistently express pluripotency markers, including SOX2 and TRA-1-80 (**Fig. 3C**). When we differentiated these hiPSCs into neural progenitor cells, we identified high proportions of dorsal telencephalic neuroepithelial markers PAX6 and FOXG1, supporting the forebrain identity of the neural progenitor cells (**Fig. 3D**). After 45 days of maturation, differentiated cortical neurons displayed a morphology with neurites and expressed mature neuron-specific markers. Staining of microtubule-associated protein 2 (MAP2) and βIII-tubulin+ was found in both soma and processes (neurites) consistent with the literature (**Fig. 3E-G**).^24, 37–39^ Moreover, we found that the vast majority of hiPSCs-derived cortical neurons expressed vesicular glutamate transporter 1 (VGLUT1), indicating an abundant presence of glutamatergic neurons (**Fig. 3E**). To determine the structural maturity of neurons, presynaptic proteins synapsin 1 and 2 (SYN1/2) were studied, which were revealed to be distributed along the axons in dotted patterns, indicating the formation of connectivity (**Fig. 3F**). Expression of cortical marker CTIP2 was also evident in culture, indicating cortical characteristics (**Fig. 3G**). Together, these data provide evidence that the CRISPR-engineered hiPSCs carrying isogenic WT and Nav1.2-L1342P can be differentiated into cortical glutamatergic neurons.

**Figure 3.**
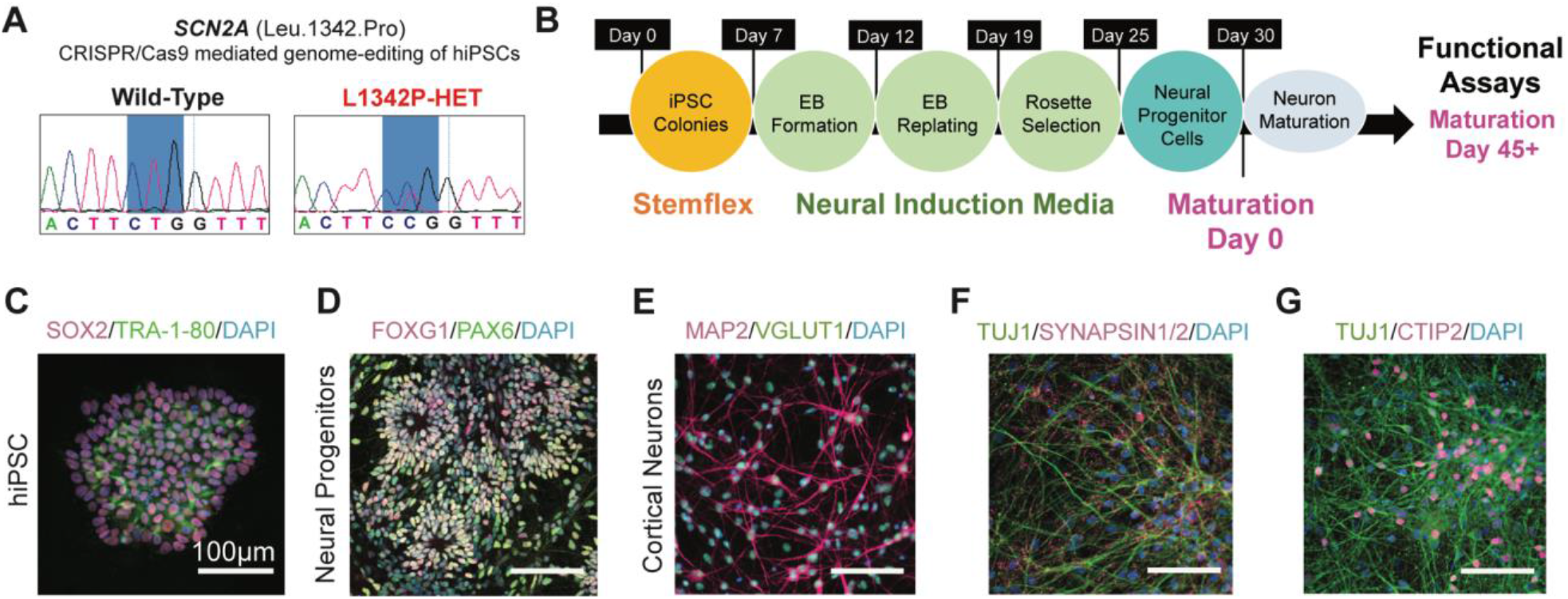
Characterization of hiPSC-derived neurons reveals a cortical neuron fate. (**A**) Sequencing of isogenic WT (black) and Nav1.2-L1342P variants (red) of hiPSCs after CRISPR/Cas9 mediated genome editing. (**B**) Schematic of a modified DUAL-SMAD differentiation protocol and the timeline for experiments. (**C**) Immunofluorescence staining of WT hiPSCs with pluripotency markers SOX2 and TRA-1-80. (**D**) Staining of neural progenitors with markers including FOXG1 and PAX6 (dorsal forebrain markers). (**E**) Immunofluorescence image of WT hiPSC-derived glutamatergic neurons expressing MAP2, colocalized with VGLUT1. (**F**) Colocalized expression of TUJ1 and synapsin1/2 was seen in the neuronal culture, suggesting that synaptic connectivity was formed among neurons. (**G**) Cortical layer marker CTIP2 indicates the cortical identity of hiPSC-derived neurons. DAPI was used to stain nuclei.

### Intrinsic excitability is enhanced in hiPSC-derived neurons carrying the Nav1.2-L1342P variant

Our results from computational modeling and whole-cell voltage-clamp recordings suggested that the neurons carrying the Nav1.2-L1342P variant would display a gain-of-function phenotype. To directly test this hypothesis, we used cortical neurons derived from hiPSCs to examine the functional consequences of the Nav1.2-L1342P variant. After maturation for at least 45 days, pyramidal-shaped neurons were selected for whole-cell current-clamp experiments (**Fig. 4A**). Neurons were held at a fixed –75 mV membrane potential, and we progressively depolarized the neurons by intracellular current injections from 0 - 135 pA, with a step of 5 pA increment. We found that the isogenic WT neurons did not fire an action potential until the injected current reached 80 pA, which was defined as the rheobase. In contrast, the L1342P neuron started to fire with a lower current injection of 60 pA (**Fig. 4B**). Quantitatively, we found a 25% reduction in rheobase of L1342P neurons, statistically different from isogenic WT neurons (**Fig. 4C**) (WT: 78.7 ± 5.3 pA, n = 27; L1342P: 57.4 ± 4.3 pA, n = 31; *p* = 0.003, *student’s t-test*). We further analyzed the voltage threshold of spiking since it is likely to be influenced by sodium channel dysfunctions ^6^. Our results revealed that L1342P neurons had a significantly lower voltage threshold, which indicates a higher probability to fire under more hyperpolarized membrane potential (**Fig. 4D**) (WT: –31.8 ± 1.5 mV, n = 27; L1342P: –36.3 ± 1.3 mV, n = 31; p = 0.02, *student’s t-test*). Furthermore, we found that the amplitude of action potential was significantly enhanced in L1342P neurons, with an average amplitude approximately 7 mV larger than that of isogenic WT (**Fig. 4E**) (WT: 20.1 ± 1.8 mV, n = 27; L1342P: 27.5 ± 1.9 mV, n = 31; *p* = 0.007, *student’s t-test*). On the other hand, we found no statistical difference in action potential (spike) width (**Fig. 4F**) (WT: 4.9 ± 0.7, n = 27; L1342P: 3.7 ± 0.3, n = 31; p = 0.37, *student’s t-test*) between the isogenic WT and L1342P neurons. In a word, our data indicate the L1342P variant makes the neurons fire action potentials more easily, making the neuron intrinsically more excitable.

**Figure 4.**
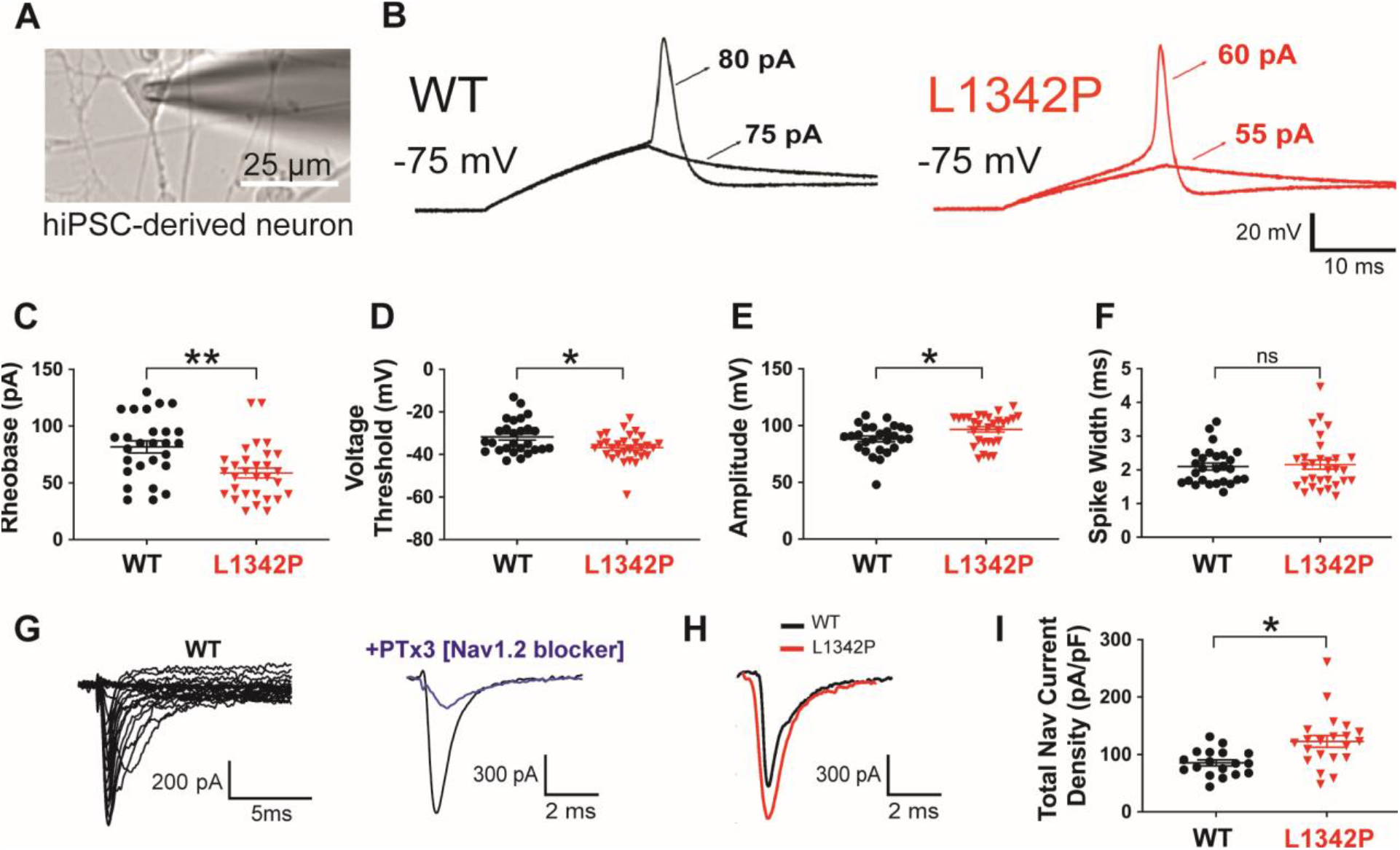
L1342P variant increases intrinsic excitability of hiPSC-derived neurons. (**A**) hiPSC-derived pyramidal-shaped neuron selected for patch-clamp experiment under the microscope. (**B**) Representative single action potential (AP) triggered by a stepwise increment of current stimulus, showing the difference in the rheobase of firing between isogenic WT and L1342P neurons at a fixed membrane potential of –75 mV. The current threshold was 80 pA for isogenic WT neurons, and 60 pA for L1342P neurons. (**C**) The L1342P variant reduced the minimal current needed for evoking an intact AP in neurons. (**D**) The voltage threshold of AP was hyperpolarized in L1342P neurons. (**E**) AP from neurons carrying the L1342P variant had elevated peak amplitude. (**F**) The L1342P variant did not change the width of the AP. (**G**) Representative voltage-dependent sodium current traces (from isogenic WT neuron, left panel), with amplitude reduction after addition of PTx3 (10 nM, right panel). (**H**) Representative sodium current comparison between isogenic WT and L1342P neurons. Traces were plotted under –20 mV depolarizing step. (**I**) Peak sodium current density was significantly increased in L1342P neurons. Data were collected from four independent differentiated batches with two clones used from each genotype. Data were analyzed by *student’s t-test*. * p < 0.05; ** p < 0.01.

Additionally, literature suggested that sodium channel variants might affect the channel trafficking and membrane expression to increase current density, contributing to neuronal hyperexcitability.^40–42^ To elucidate whether the L1342P variant may affect neuronal excitability in a similar manner, we recorded sodium current from our hiPSC-derived neurons in voltage-clamping mode. We were able to record a family of voltage-dependent inward current traces from the hiPSC-derived neurons carrying WT and L1342P mutant channels, with peak current nearly reaching 1 nA (**Fig. 4G left**). After applying phrixotoxin-3 (PTx3, 10 nM), a specific Nav1.2 channel blocker at or below 30 nM,^43–45^ the peak current was significantly reduced (**Fig. 4G right**). This result suggests that Nav1.2 current is one of the major components contributing to the total sodium current in these neurons, consistent with the literature.^6^ When we compared the peak sodium current density between isogenic WT and L1342P neurons, we found a significantly greater sodium current density in L1342P neurons (**Fig. 4H-I**) (WT: 85.5 ± 5.3 pF/pA, n = 18; L1342P: 122.9 ± 10.3 pF/pA, n = 21; p = 0.003, *student’s t-test*). A higher current density of sodium channel, together with reduced rheobase and voltage threshold, are likely to serve as the basis underlying the increased intrinsic excitability of neurons carrying the L1342P variant.

### Repetitive firing is elevated in hiPSC-derived neurons carrying the Nav1.2-L1342P variant

To further investigate how the L1342P variant affects repetitive action potential firings, we performed whole-cell current-clamp recording using a prolonged 400 ms current stimulus ranging from 0 to 125 pA in 5 pA increments. From the 45 to 55 pA stimulus, L1342P neuron fired 2-4 action potentials in response to current steps (**Fig. 5A**). In contrast, single or at most two action potentials were elicited in isogenic WT neurons (**Fig. 5A**). Overall, the L1342P neurons fired significantly more action potentials than isogenic WT in the range of 0 - 125 pA (repeated-measures two-way *ANOVA* analysis, F (1, 39) = 6.67, p = 0.014) (**Fig. 5B**). It is worth noting that we also observed a distinct pattern from the input-output relationship of isogenic WT and L1342P neurons. Under depolarizing current within 0 - 20 pA, the firing frequency elicited between two groups in each step is similar. Starting from 20 pA, we observed a steeper slope of increase in firing from the L1342P neurons until 75 pA current injection, indicating that they are more responsive to stimulus during this range. However, the curve from L1342P neurons fell between the range of 75 - 125 pA, while isogenic WT neurons retained a steady firing frequency after achieving the plateau. Importantly, the maximum number of action potential firing that can be triggered by current injection was significantly higher in L1342P neurons, showing a 35% increase over WT neurons (**Fig. 5C**) (WT: 3.4 ± 0.3, n = 18; L1342P: 4.6 ± 0.4, n = 23; p = 0.02, *student’s t-test*). Together, our data suggest the L1342P variant greatly influences the repetitive firing, leading to the hyperexcitability phenotype of neurons.

**Figure 5.**
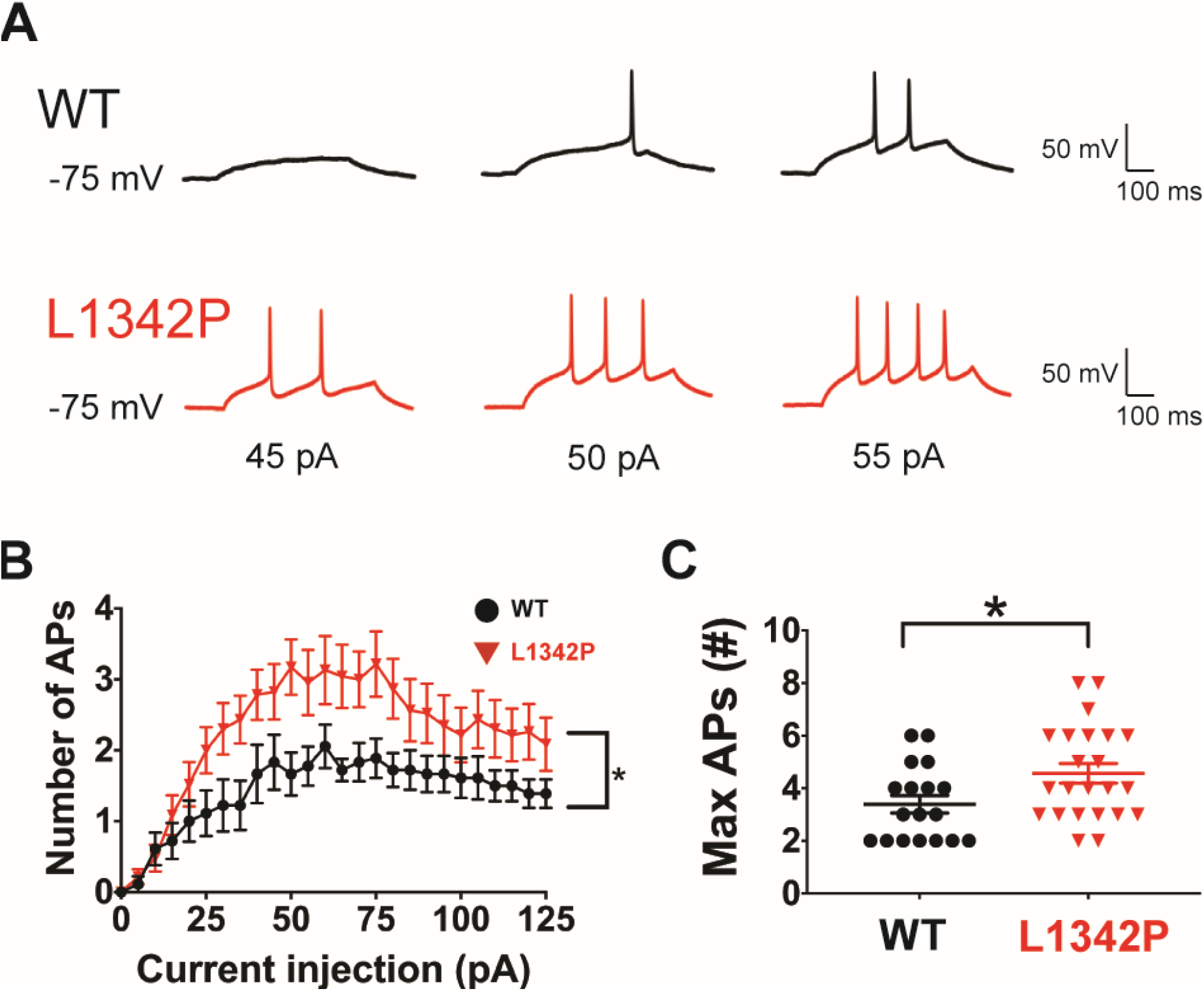
L1342P variant enhances the repetitive firing of hiPSC-derived neurons. (**A**) Representative sustained action potential firings from hiPSC-derived cortical neurons expressing isogenic WT (black) and L1342P (red) mutated Nav1.2. (**B**) AP number per epoch in response to graded inputs from 0 to 125 pA current injection is shown. L1342P neurons constantly fired more APs than isogenic WT neurons. (**C**) Maximum number of APs triggered from each neuron under the range of 0-125 pA current injections. Data were collected from four independent differentiated batches, with two clones used from each genotype. Data were analyzed by *student’s t-test.* * p < 0.05.

### Neural network excitability is heightened in hiPSC-derived neurons carrying the Nav1.2-L1342P variant

We observed elevated evoked action potential firing of individual neurons carrying the Nav1.2-L1342P variant in our single-cell patch-clamp recording. However, whether this enhanced intrinsic excitability translates into higher activities in a neural network is unknown. MEA is a relatively high-throughput extracellular recording assay that can record the firing of a population of neurons in their normal culture medium, providing a physiologically relevant avenue to assess neuronal network excitability (**Fig. 6A**).^22, 46, 47^

**Figure 6.**
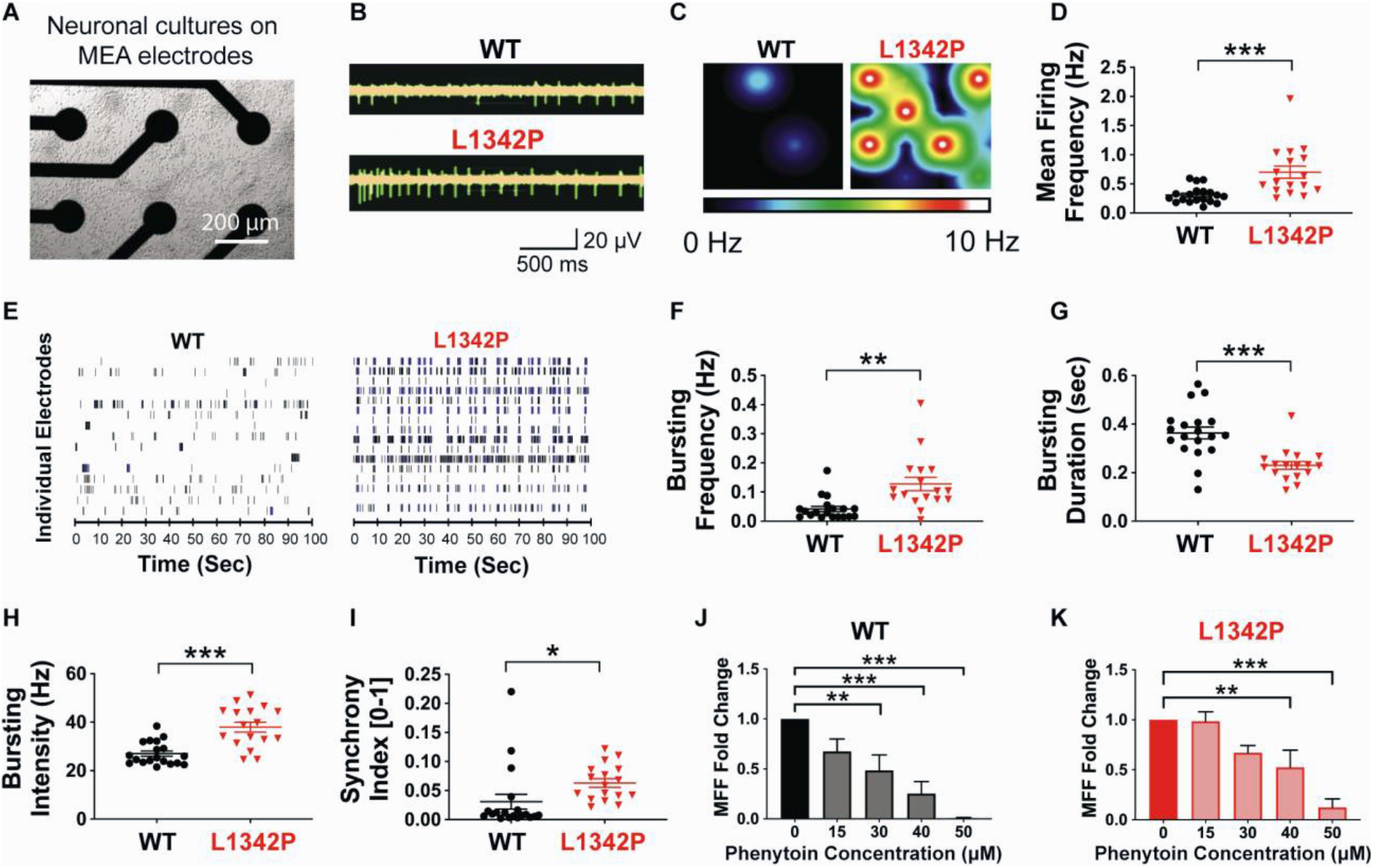
Micro-electrode array (MEA) recording of the firing of hiPSC-derived neurons carrying WT and L1342P variant. (**A**) MEA well plated with hiPSC-derived neurons. (**B**) Representative raw spikes from isogenic WT and L1342P cultures. Each spike indicated a spontaneous activity. (**C**) Heat-maps recordings of neuronal firing. The intensity of firing frequency is color-coded, with warm colors (white, red, orange, and yellow) indicating high firing frequency and cool colors (green and blue) depicting low firing frequency. Each color circle represents an active electrode. (**D**) Mean firing frequency (MFF) comparison between isogenic WT and L1342P culture. (**E**) Representative spike raster plots from isogenic WT or Nav1.2-L1342P culture. Each row represents the spikes recorded from one electrode through 100 seconds, with each tick indicating one spontaneous event. Bursting events are depicted by a cluster of ticks in blue. (**F-I**) Parameters quantitatively describing the network activities in neuron culture, including bursting frequency (F), bursting duration (G), and bursting intensity (H), Synchrony Index (I). Data were pooled from wells across four hiPSC lines (two for each genotype), with n = 19 (wells) for isogenic WT and n = 17 (wells) for the L1342P culture. The *student’s t-test* was performed when data is normally distributed (Kolmogorov-Smirnov Normality Test), and Mann-Whitney U was performed when data is not normally distributed. * p < 0.05; ** p < 0.01. (**J-K**) Inhibitory effects of different dosages of phenytoin between isogenic WT (J) and L1342P cultures (K). Data are presented as a fold change of mean firing frequency in response to increasing concentrations of phenytoin from 15-50 μM. The sample size was n = 7, 8, 6, 6 (wells) for different concentration in WT culture, and n = 6, 6, 6, 8 (wells) for L1342P culture. One-way ANOVA was performed, with Bonferroni corrections compared to the well before adding the phenytoin. * p < 0.05; ** p < 0.01; ***p < 0.001.

Spontaneous neuronal network activity was recorded using MEA, which showed that the L1342P neuron culture fired at a higher level than the isogenic WT culture in traces of a 2.5-sec epoch (**Fig. 6B**). The firing of a population of neurons could be visualized in a heat-map view. Each colored circle represented an active electrode (**Fig. 6C**), with blue suggesting low firing frequency and yellow/red indicating high firing frequency. In a representative isogenic WT culture, two active electrodes exhibited low activity, whereas the L1342P culture had five active electrodes with higher activity (**Fig. 6C**). Quantitatively, MEA recording revealed that the mean firing frequency of L1342P culture was significantly elevated compared with isogenic WT (**Fig. 6D**) (WT: 0.31 ± 0.03 Hz, n = 19; L1342P: 0.70 ± 0.1 Hz, n = 17; p < 0.001, *student’s t-test*). To further understand the temporal behavior of these neurons, we analyzed the raster plot, in which the recorded activity of each electrode was plotted over time. While we observed a similar number of active electrodes across wells, we detected more bursting events in the L1342P neuron culture compared to the isogenic WT culture (**Fig. 6E**). In the isogenic WT culture, the active electrodes were firing at a relatively low frequency with scattered firing characteristics over 100 seconds. In contrast, the representative well-wide raster plot of L1342P culture showed a higher firing rate with notable bursting events over the same recording window (**Fig. 6E**).

Bursting events, including the bursting frequency, bursting duration, and bursting intensity (spiking frequency within each bursting event), were further quantified as these are seizure-related firing characteristics of the neuronal culture.^36, 48–50^ The bursting frequency was significantly increased in the L1342P culture compared to isogenic WT (**Fig. 6F**) (WT: 0.04 ± 0.009, n = 19; L1342P: 0.13 ± 0.02, n = 17; p = 0.0002, Mann-Whitney U). Interestingly, while the bursting duration was shortened significantly in L1342P culture (WT: 0.36 ± 0.02, n = 19; L1342P: 0.23 ± 0.01, n = 17; p < 0.0001, *student’s t-test*) (**Fig. 6G**), the bursting intensity was markedly elevated, which was reflected by the decreased inter-spike interval (ISI) within the burst (**Fig. 6H**) (WT: 27 ± 1.1 Hz, n = 19; L1342P: 38 ± 2.0 Hz, n = 17; p < 0.0001, *student t-test*). Furthermore, we measured the synchrony index, which was determined by synchrony of spike firing between electrode pairs,^29^ to reflect the strength of synchronized activities between neurons (value is between 0-1, with 1 indicating the highest synchrony). The synchrony index was markedly increased in L1342P neuron culture (**Fig. 6I**), suggesting seizures-related hypersynchronization (WT: 0.03 ± 0.01, n = 19; L1342P: 0.06 ± 0.006, n = 17; p = 0.04, Mann-Whitney *U*). These results are consistent with the observation of enhanced intrinsic excitability of L1342P neuron culture in patch-clamping recording, while further demonstrate elevated network excitability, including bursting and synchronous firing caused by the L1342P variant.

### Sensitivity towards anticonvulsant phenytoin is reduced in hiPSC-derived neurons carrying the Nav1.2-L1342P variant

In clinical observations, patients carrying the Nav1.2-L1342P variant develop intractable seizures with a minimal response towards commonly used antiepileptic drugs.^16^ To test if we could recapitulate elements of this clinical observation in a cell-based model, we studied a commonly used first-line anticonvulsant phenytoin using the MEA platform. Phenytoin is a general sodium channel blocker and has been shown to inhibit high-frequency repetitive spikings to control seizure episodes.^51–54^ We found that in isogenic WT culture, 15 μM phenytoin started to reduce the mean firing rate, and a statistically significant reduction was reached with phenytoin at 30 μM (**Fig. 6J**). However, 15 μM phenytoin did not reduce the mean firing rate of L1342P culture at all, and 40 μM of phenytoin was required to achieve a statistically significant reduction of mean firing rate in L1342P culture (**Fig. 6K**). Our data thus suggests that neurons carrying the L1342P variant are likely to have a reduced sensitivity towards phenytoin, providing a plausible explanation for the clinical observation that patients carrying the Nav1.2-L1342P variant are largely resistant to the standard therapies.

### Nav1.2 specific inhibitor PTx3 reduces the spontaneous and chemically-induced firing of hiPSC-derived neurons carrying the Nav1.2-L1342P variant

General sodium channel blockers (e.g., phenytoin) have limited efficacy for many patients with a variety of different sodium channel variants.^5, 51, 52, 55^ Therefore, the development of isoform-specific blockers to treat patients carrying sodium channel variants in a precision medicine manner is desirable. Although Nav1.6 specific or Nav1.6/Nav1.2 dual blockers have been reported,^56^ isoform-specific blockers for Nav1.2 are not currently available. Neurotoxin phrixotoxin-3 (PTx3) has been suggested as a Nav1.2 specific blocker when used at a dose at or below 30 nM.^43–45^ While PTx3, as a neurotoxin, may not be directly useful in a clinical setting, to provide proof-of-concept evidence of the utility of Nav1.2 specific blocker for clinical translation, we studied the inhibitory effect of PTx3 on our neuronal cultures.

We used MEA to test the inhibition effect of PTx3 on baseline neuronal firings. After adding PTx3, the neuronal firing from L1342P cultures was reduced markedly (**Fig. 7A**). Quantitatively, a highly effective and significant reduction of baseline firing was observed in both isogenic WT and L1342P cultures (**Fig. 7B**) (WT: 0.39 ± 0.1, n = 6; p = 0.03. L1342P: 0.32 ± 0.01, n = 6; p = 0.03, Wilcoxon test, compared to the untreated baseline). Additionally, we evaluated the potency of PTx3 to inhibit neuronal firings under the condition of a chemically induced hyperexcitability to further model seizure states. Kainic acid (KA, 5 μM), a glutamate receptor agonist, also a commonly used compound to chemically trigger epileptiform activities, was used.^57, 58^ KA was able to markedly enhance the firings of both isogenic WT and L1342P culture (**Fig. 7C top**). With the addition of PTx3, we observed a strong reduction in the spike numbers (**Fig. 7C bottom)**. Both mean firing frequency and bursting frequency were significantly reduced by PTx3 in both isogenic WT and L1342P neuronal culture (**Fig. 7D-E**) (MFF: WT: 0.26 ± 0.07, n=8, p = 0.007; L1342P: 0.20 ± 0.07, n = 6, p = 0.03; bursting frequency: WT: 0.27 ± 0.1, n=8, p = 0.007; L1342P: 0.07 ± 0.06, n = 6, p = 0.03; Wilcoxon test, compared to the KA-induced activity). Interestingly, PTx3 seems to have a slightly stronger inhibition effect on L1342P culture compared to those on isogenic WT culture (**Fig. 7D-E**). Our results show that the PTx3 could effectively alleviate the hyperexcitability of neuronal culture with the L1342P variant, supporting the hypothesis that isoform-specific Nav1.2 blockers could potentially be novel anticonvulsant drugs.

**Figure 7.**
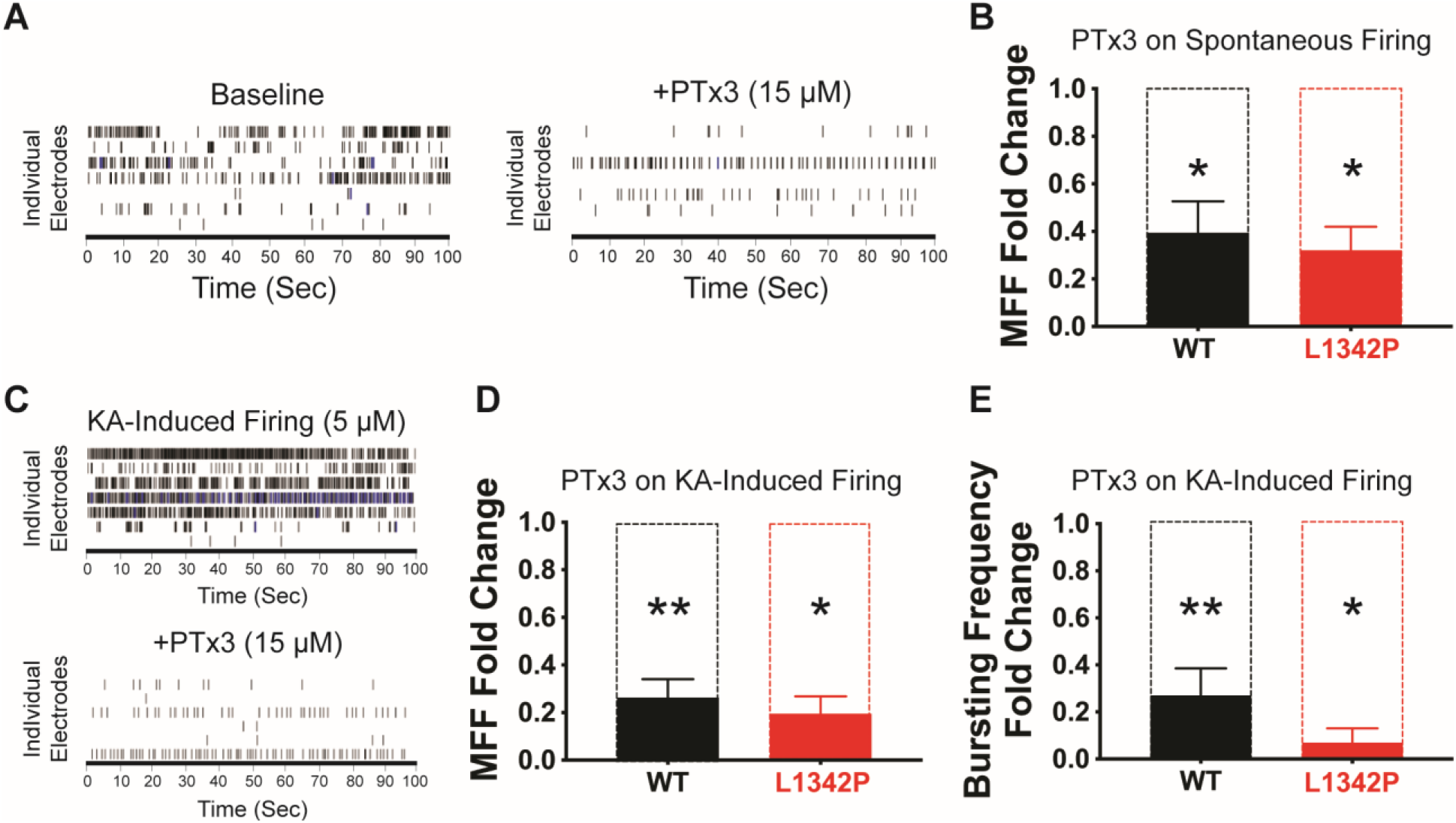
Phrixotoxin-3 (PTx3) effectively suppresses spontaneous and kainic acid (KA)-induced firings of hiPSC-derived neurons carrying the L1342P variant. (**A**) MEA recording showed that baseline mean firing frequency was reduced after the addition of PTx3 in L1342P culture. (**B**) PTx3 significantly inhibited the baseline firing in both isogenic WT and L1342P culture. Wilcoxon test was performed to compare the neuronal firing before and after the PTx3 treatment. * p < 0.05. (**C**) KA enhanced neuronal firing by elevating the bursting, which can be effectively inhibited by PTx3. (**D**) PTx3 significantly inhibited the mean firing frequency in both WT and L13432P neuron culture after KA-stimulation. (**E**) PTx3 significantly inhibited the bursting frequency in both WT and L13432P neuron culture after KA-stimulation. For baseline firing reduction analysis, n = 6 (wells) for WT, and n = 6 (wells) for L1342P culture. For inhibiting the KA-induced firing using the PTx3, n = 8 (wells) for WT, and n = 6 (wells) for L1342P culture. Wilcoxon test was performed to compare the KA-treated cultures before and after the PTx3 treatment. * p < 0.05; ** p < 0.01.

## Discussion

In this study, we revealed substantial and complex biophysical property changes in the L1342P variant, which resulted in hyperexcitability of cortical neurons. In particular, we found that cortical neurons derived from hiPSCs carrying heterozygous L1342P display significantly enhanced sustainable action potential firings, larger spike amplitude, hyperpolarized voltage threshold, lower rheobase, and greater sodium current density. By investigating the network activity of neuronal populations using MEA, we revealed increased network excitability with burst firing and a higher synchrony index in the L1342P culture. Remarkably, we obtained evidence to show that hiPSC-derived neurons carrying the L1342P variant are less sensitive to a commonly used anticonvulsant compound phenytoin, but have equal or slightly stronger sensitivity to a Nav1.2 isoform-specific blocker.

Nav1.1, Nav1.2, and Nav1.6 are three major voltage-gated sodium channels expressed in the central nervous system.^59^ Notably, variants in all these channels are associated with seizures, probably due to their important roles in mediating action potential firing.^60, 61^ Among them, Nav1.1 is strongly expressed in the interneurons.^62^ Majority of the seizure-associated Nav1.1 variants are suggested to be loss-of-function, which reduces the excitability of inhibitory interneurons, leading to less inhibition on principal excitatory neurons.^63^ In turn, without sufficient inhibition, excitatory neurons become more excitable and promote seizures. On the other hand, both Nav1.2 and Nav1.6 are predominantly expressed in the principal excitatory neurons.^6^ Epilepsy-associated Nav1.6 variants are mainly gain-of-function. Studies found that enhanced excitability of hiPSC-derived excitatory neurons is the primary mechanism underlying the gain-of-function Nav1.6 variant related seizure.^64^ Loss-of-function variants of Nav1.6 are also found in patients, but these patients did not develop epileptic phenotypes.^65^ In contrast, the association between variants in Nav1.2 and seizure are more complicated. Intriguingly, seizures have been observed in both gain-of-function and loss-of-function Nav1.2 variants.^5, 66^ Since Nav1.2 is expressed in the principal excitatory neurons, it is predicted that Nav1.2 variants would enhance neuronal excitability. Nevertheless, this hypothesis has not been explicitly tested in human neurons. Here we showed, for the first time, that the epilepsy-associated Nav1.2 variant L1342P causes hyperexcitability and hypersynchronous firing in human excitatory neurons derived from hiPSCs. Mechanistically, we revealed that the L1342P variant resulted in an increased window current and enhanced current density.^35, 50^ We proposed a working hypothesis that the extra Nav1.2 channels available in the cell membrane, which possess altered biophysical properties to be more readily activated, is likely to push neurons into a hyperexcitable state to promote seizures.

Interestingly, despite an overall gain-of-function phenotype associated with neuronal hyperexcitability, we also observed traits of loss-of-function, particularly in action potential firing. In the input-output relationship, we found the L1342P neurons initially had a strong response to the current stimulus by increasing AP firing significantly. However, the L1342P neurons later failed to maintain sustainability in firing frequency at high current injection (**Fig. 5B**). A similar trend of the input-output relationship was reported in a previously described variant Q1478K in Nav1.1.^67^ The Q1478K variant caused a “self-limiting” hyperexcitability phenotype, possibly caused by a significant reduction in sodium current density. ^67^ However, we found an increased current density in L1342P, suggesting a different mechanism. We propose that the failure to maintain spiking could be a result of enhanced inactivation of the Nav1.2-L1342P mutant channel, which was discovered in our biophysical assays. Further studies are required to test this proposed mechanism.

Several recently reported variants in *SCN2A* from patients with childhood epilepsy also presented with evidence indicating developmental impairments.^68^ The terminology of “developmental and/or epileptic encephalopathy” thus has been used to imply that the developmental impairments could occur independently of seizures.^69, 70^ Indeed, profiling of patients with the L1342P variant revealed developmental disorders.^13, 14, 16^ Three out of five L1342P cases display microcephaly, brain volume loss, and reduced head circumference. It is currently not known whether the developmental impairments observed in patients carrying L1342P are related or independent of the seizures caused by the L1342P variant. These disease symptoms related to neurodevelopment still need to be carefully investigated in the future.

Studies have traditionally used the heterologous expression system as a convenient platform to study ion channel variants related to epilepsies. This type of analysis is informative regarding the specific biophysical properties of a particular variant.^28, 31, 71^ Moreover, this biophysical analysis can also provide information regarding “gain-of-function” versus the “loss-of-function” feature of a particular variant of interest.^40, 72^ Determining the gain versus loss-of-function is critically important, both for establishing genotype-phenotype correlations and informing clinical practice. Indeed, an insightful clinical study has shown that seizures patients carrying traditionally “gain-of-function” Nav1.2 variants can be adequately controlled by sodium channel blockers, whereas this same class of sodium channel blockers exacerbate seizures and worsen outcomes for patients carrying loss-of-function Nav1.2 variants.^5, 66^ Using electrophysiology and human neurons derived from hiPSCs, here we provided evidence to show that neuron carrying Nav1.2-L1342P variant is clearly gain-of-function, despite complex biophysical properties revealed in the heterologous expression system.

Even with advanced genetic testing, controlling seizures is still challenging in clinical practice. The standard antiepileptic drug (AED) guidelines are usually applied to patients indiscriminately.^12, 73^ A large number of patients do not respond to the first antiepileptic drug that was prescribed to them and they have to go through a trial-and-error process with different medication. This tedious and often ineffective process significantly increases disease burden and may cause the optimal treatment window to be missed for children with epilepsy. Therefore, there is an urgent need to explore new approaches to guide clinical practice in a precision medicine manner. hiPSCs-based *in vitro* genetic models hold great promise to transform clinical practice. Neurons derived from hiPSCs are amenable for high-throughput parallel screening for targeted drug discovery. Pairing with MEA, a high-throughput, extracellular recording assay of neuronal activity,^57, 74, 75^ such studies have been attempted with neurons carrying Nav1.6 variants.^64^ Using the hiPSC-derived neurons, it was recently found that riluzole and phenytoin can target specific Nav1.6 variants to attenuate bursting phenotypes *in vitro*. Importantly, some effective compounds identified in the hiPSC model successfully suppress seizure events of patients in the clinical study, highlighting the translational value of the *in vitro* hiPSC-based platform.^64^

While the use of patient-derived hiPSC line is desirable and could directly model subject-specific phenotypes and interindividual differences,^47^ reference hiPSCs with genetic variants engineered by CRISPR/Cas9 is an indispensable complement approach. This approach eliminates the need to obtain samples from patients, standardize the genetic background for cross variants comparison, and provides a reproducible baseline control for potential drug discovery targeting recurring genetic variants. Here in this study, we showed that reference hiPSC-derived neurons carrying the drug-resistant Nav1.2-L1342P variant were not sensitive to phenytoin but had equal or even stronger sensitivity towards an experimental isoform-specific Nav1.2 blocker. Our results demonstrated the usefulness of such an *in vitro* platform to potentially advance drug selection and discovery, which warrants additional studies to streamline this process for future clinical translation.

In summary, we revealed a hyperexcitability phenotype in hiPSC-derived neurons carrying the Nav1.2-L1342P variant, with unique features in altering the neuronal excitability as possible pathogenic mechanisms. Moreover, we demonstrated that our *in vitro* model is able to partially recapitulate the pharmaco-responsiveness of the Nav1.2-L1342P variant. Using this *in vitro* model, we further show, as a proof-of-principle, that isoform-specific Nav1.2 blocker may be an effective treatment of hyperexcitability caused by Nav1.2 gain-of-function variants. Our results highlight the utility of genome-edited, hiPSC-derived neurons for mechanistic investigation of disease-causing variants, as well as for drug discovery to advance personalized treatment for patients carrying a variety of genetic variants.

## Acknowledgments

We thank Dr. Amy Brewster for the critical reading of this paper.

## Funding

The authors gratefully acknowledge support from the Purdue University Institute for Drug Discovery and Institute for Integrative Neuroscience. Yang lab is grateful to the *FamilieSCN2A* foundation for the Action Potential Grant support. Y.Y. is supported by funds from Ralph W. and Grace M. Showalter Research Trust, and Purdue Big Idea Challenge 2.0 on Autism. M.I.O.A. is supported by the Fulbright-Colciencias scholarship program. M.E. is supported by the National Science Foundation (NSF) Graduate Research Fellowship Program (GRFP) (DGE-1842166). This project was also funded, in part, with support from the Indiana Clinical and Translational Sciences Institute, in part by Award Number UL1TR002529 from the National Institutes of Health, National Center for Advancing Translational Sciences, Clinical and Translational Sciences Award. The content is solely the responsibility of the authors and does not necessarily represent the official views of the National Institutes of Health or the National Science Foundation.

## Competing Interests

The authors report no competing interests.

## Author contributions

Z.Q. and Y.Y. designed experiments; Z.Q. and M.I.O.A. performed research; Z.Q., M.I.O.A., and Y.Y. analyzed data; J.Z., M.E., A.M.T., X. C., J. M. S., L. Y., and R. A. O. participated in experiments and data analysis. J.A.S., J.C.R., A.B.B, C.Y, Z.H., W.C.S, participated in data analysis and research design. Z.Q. and Y.Y. wrote the paper with inputs from all authors.

## Notes

### Competing Interest Statement

The authors have declared no competing interest.

